# A new genetic method for diet determination from faeces that provides species level resolution in the koala

**DOI:** 10.1101/2023.02.12.528172

**Authors:** Michaela D. J. Blyton, Kylie L. Brice, Katarzyna Heller-Uszynska, Jack Pascoe, Damian Jaccoud, Kellie A. Leigh, Ben D. Moore

## Abstract

An animal’s diet is a crucial trait that defines their realised ecological niche, especially for dietary specialists such as the koala (*Phascolarctos cinereus*), a threatened arboreal marsupial folivore. Unfortunately, the current methods used to characterise koala diet are labour intensive, biased and/or unreliable. Further, in this study we show that four barcoding genes (*ITS, ETS, CCR* and *matK*) are unable to resolve potential koala food trees to species. Therefore, we developed and tested a novel SNP-based method for the analysis of koala diet from faeces using the DArTseq platform. This method returned a large number of species-specific SNPs for candidate koala food tree species. Due to low within-species variation, few individuals of each tree species are needed to capture the majority of DArTseq SNP diversity. Nonetheless, we suggest sampling multiple trees to reduce the impact of high allele dropout rates in the DArTseq data. After identifying species-specific SNPs from candidate food tree species from two study sites with different assemblages of eucalypts we were able to detect those SNPs in koala faecal DNA using DArTag, a targeted genotyping assay. This enabled us to semi-quantitatively characterise the koalas’ diets. The food tree species identified were in broad agreement with previously known koala food tree species but also revealed additional species that may contribute to koala diet. This approach provides an important new tool for use in koala ecology and conservation and may prove useful in diet determination for other species where high taxonomic resolution is crucial and dietary DNA is scarce.

## Introduction

What animals eat underpins the nutrition, health and reproduction of individuals and the growth and persistence of wild animal populations. Diet choice also structures food webs, determines the flow of energy and nutrients through ecosystems and shapes ecological communities. Obligatory specialist feeders (Shipley, Forbey, & Moore, 2009) exhibit particularly narrow dietary niches, making the accurate identification of their small number of dietary species essential to assessing and mapping habitat quality. Additionally, distinguishing closely related food species can be important for generalist feeders, because nutritional quality can differ substantially between congeneric species (e.g. Craig, Bell, & Atkins, 1991)

The koala (*Phascolarctos cinereus*) is an arboreal marsupial folivore that is now threatened with extinction throughout the central and northern parts of its range (Gonzalez-Astudillo, Allavena, McKinnon, Larkin, & Henning, 2017; Gordon, Hrdina, & Patterson, 2006; McAlpine et al., 2015). Conservation efforts for the species have focused on the protection of koala habitat, with the presence of preferred food trees recognised in legislation as the key attribute of suitable areas (Adams-Hosking, McAlpine, Rhodes, Grantham, & Moss, 2012; Lunney, Phillips, Callaghan, & Coburn, 1998; White, 1999). It is well understood that the koala feeds almost exclusively on the foliage of eucalypts (trees from the genera *Eucalyptus*, *Angophora* and *Corymbia*). However, it is more difficult to confidently determine which eucalypt species are actually eaten by koalas and to what extent (Moore & Foley, 2000). Feeding is often inferred from faecal pellet deposition or diurnal patterns of tree occupancy, which can be misleading, as koalas choose trees for shelter and social reasons in addition to food quality, and do not feed in all trees they use, particularly during the day (Ellis, Melzer, Carrick, & Hasegawa, 2002; Marsh, Moore, Wallis, & Foley, 2014a).

Techniques for characterising herbivore diets have been comprehensively reviewed elsewhere (e.g. Mayes and Dove, 2000; Garnick et al 2018) and include direct behavioural observation and the analysis of postingestive samples (e.g. stomach contents or faeces) using microhistology, near infrared reflectance spectroscopy, stable isotope analysis, chemical analysis of plant cuticular wax alkanes and DNA barcoding. Direct observation or acoustic recording of time spent feeding is very rarely used with koalas due to their labour intensive nature (e.g. Logan & Sanson, 2003; Marsh, Moore, Wallis, & Foley, 2014b). Microhistological analysis is an established technique, in which leaf cuticle fragments are matched to a reference collection of candidate food trees (Castle et al., 2020; Goldberg et al., 2020; King & Schoenecker, 2019), and has been applied to koalas in numerous studies (e.g. Melzer, Cristescu, Ellis, FitzGibbon, & Manno, 2014; Nyo Tun, 1994; Wu, McAlpine, & Seabrook, 2012). The technique is labour intensive, requires specific expertise, is potentially subject to bias, and we have found that it is unable to distinguish all species of *Eucalyptus* in some areas of the koala’s range. The faecal *n*-alkane composition originating from cuticular waxes of consumed plants is another measure that has been used to assess diet composition in koalas and other herbivores where there are few potential dietary species (Brice et al., 2019; Dove & Mayes, 2005). However, this method can only distinguish as many diet items as there are markers available and thus is not effective for diets with many components, or where wax profiles do not differ much between species (Bugalho, Dove, Kelman, Wood, & Mayes, 2004). Given the shortcomings of these methods, we aimed to assess whether techniques of diet composition analysis based upon the molecular analysis of faecal samples could offer a practical and reliable solution to koala researchers and managers.

Traditionally, genetic determination of diet composition has been performed by amplifying conserved ‘barcoding’ genes from the faeces and matching the recovered sequences to a representative database. This approach has proven highly successful in cases, where the candidate food species are taxonomically diverse (Castle et al., 2020; Goldberg et al., 2020; Kartzinel et al., 2015; King & Schoenecker, 2019). However, eucalypts can be difficult to differentiate genetically due to incomplete lineage sorting, frequent hybridisation and unresolved phylogenetic relationships (Jones, Nicolle, Steane, Vaillancourt, & Potts, 2016; Schuster et al., 2018; Thornhill et al., 2019). As such, several commonly used barcoding genes have been unable to resolve some closely related eucalypt species (Buys et al., 2016; Fladung, Schroeder, Wehenkel, & Kersten, 2015). Nonetheless, to test the utility of this well-established approach for the determination of koala diets, we assessed five barcoding genes (*ITS, ETS, matK, ndh* and *CCR*) that have previously been used in *Eucalyptus* phylogenetic analyses (Gadek, Wilson, & Quinn, 1996; Lucas et al., 2007; Poke, Martin, Steane, Vaillancourt, & Reid, 2006; Thornhill, Ho, Külheim, & Crisp, 2015).

Recently, next-generation sequencing (NGS) of single nucleotide polymorphisms (SNPs) have been found to reliably differentiate closely related species within several plant genera, including *Eucalyptus* (Dasgupta, Dharanishanthi, Agarwal, & Krutovsky, 2015; Ndjiondjop et al., 2018; Yuskianti, Fa Xin, Bian Xiang, & Shiraishi, 2011). SNPs that act as species-specific markers may also be useful in analysing diet composition from faecal samples and may provide greater taxonomic resolution than barcoding genes. One SNP-based method is DArTseq (Diversity Arrays Technology Sequencing), which sequences a reduced fraction of the genome that corresponds to predominantly active genes. This selection is achieved through the use of a combination of restriction enzymes which separate low copy sequences from the repetitive fraction of the genome (Jaccoud, Peng, Feinstein, & Kilian, 2001; Kilian et al., 2012). We used this platform to identify species-specific SNPs from a reference set of candidate koala food trees from two study sites and then quantified those SNPs in faecal DNA. In assessing this new approach to diet determination, we also examined the sampling requirements for the candidate dietary species in order to adequately capture their genetic diversity.

The potential for DArTseq to detect *Eucalyptus* SNPs from koala faeces has been demonstrated previously (Schultz, Cristescu, Littleford-Colquhoun, Jaccoud, & Frère, 2018). However, faecal DNA is derived primarily from microbes and subsequently only a very small fraction is diet-associated DNA. As a result, non-selective sequencing of faecal DNA is likely to return a low coverage of the target sequences (Srivathsan, Ang, Vogler, & Meier, 2016). Therefore, to improve our detection of dietary species-specific SNPs in faecal DNA we trialled DArTag, a targeted genotyping assay, which is already broadly adopted for breeding, genetic analysis and monitoring applications in plants and animals (https://www.diversityarrays.com/technology-and-resources/targeted-genotyping/). This method uses custom designed oligos to amplify targeted SNPs, and their flanking sequences (∼ 50 bp), prior to NGS sequencing. In this way the representation of target sequences is increased. Here we report the performance of a DArTag panel developed based on the DArTseq data to selectively amplify the eucalypt SNPs from the scats.

Through our analysis and development of these new genetic methods for determining koala dietary tree species from scats we determine the utility of these approaches for use in koala ecology, management and conservation. Additionally, as the first application of DarTseq and DarTag to faecal diet determination, this study highlights the potential for these methods to be used for diet determination of other species where high taxonomic resolution is crucial and dietary DNA concentration in faeces is poor.

## Materials and Methods

### Study sites and sample collection

#### Captive Koalas

Faeces were sampled from six wild-caught koalas from Cape Otway, Victoria, Australia, held temporarily in captivity and fed exclusively on either of two *Eucalyptus* species (*E. viminalis:* n=3; *E. obliqua* n=3). Details of husbandry can be found in Blyton et al. (2019).

#### Mountain Lagoon, Blue Mountains, New South Wales

Koalas have a patchy distribution in the Blue Mountains (New South Wales, Australia) (Lunney, Crowther, Shannon, & Bryant, 2009), that likely reflects the heterogeneous mix of vegetation communities in the region. Our study investigated the diet of a koala population at Mountain Lagoon (-33.4442 S, 150.6295 E), that occupies a variety of vegetation communities including: Sydney hinterland sheltered turpentine–apple forest, Sydney hinterland peppermint–apple forest, Lower Blue Mountains exposed red bloodwood forest, Blue Mountains grey gum–stringybark transition forest, Blue Mountains blue gum–turpentine gully forest and Blue Mountains shale cap forest (Gallahar et al. 2021). Koalas at the site had access to eleven eucalypt and one non-eucalypt tree species in the family Myrtaceae from which they could have fed (Table 1).

**Table 1:** Species-specific SNPs identified in potential dietary species at Mountain Lagoon.

| Genus (subgenus) | Common Name<br>(Species) | Fixed private<br>SNPs (strict) | Subset for<br>Diet Analysis<br>(strict) | Most Suitable<br>for DarTag<br>(strict) | Designed<br>Tags<br>(strict) |
| --- | --- | --- | --- | --- | --- |
| *Eucalyptus*<br>(*Eucalyptus* *) | Sydney Peppermint<br>(*E. piperita*) | 618 (458) | 614 (455) | 140 (105) | 15 (15) |
|  | Blue-Leaved<br>Stringybark<br>(*E. agglomerata*) | 503 (349) | 499 (346) | 128 (86) | 3 (3) |
|  | White Stringybark<br>(*E. globoidea*) | 510 (362) | 501 (359) | 122 (89) | 15 (15) |
| *Eucalyptus*<br>(*Symphyomyrtus*) | Mountain Grey Gum<br>(*E. cypellocarpa*)<sup>#</sup> | 696 (9) | 91 (9) | 18 (1) | 10 (2) |
|  | Mountain Blue Gum<br>(*E. deanei*) | 1255 (85) | 430 (85) | 91 (19) | 16 (16) |
|  | Grey Gum<br>(*E. punctata*) | 2746 (474) | 1243 (466) | 254 (95) | 67 (67) |
|  | Sydney Blue Gum<br>(*E. saligna*) | 1857 (968) | 1809 (950) | 397 (201) | 16 (16) |
|  | Grey Ironbark<br>(*E. paniculata*) | 1138 (744) | 1088 (711) | 226 (147) | 14 (14) |
|  | Narrow-leaved<br>Ironbark (*E.*<br>*beyeriana*) | 2727 (10) | 350 (10) | 66 (1) | 7 (2) |
| *Corymbia* | Red Bloodwood<br>(*C. gummifera*) | 1124 (908) | 1117 (905) | 208 (157) | 13 (13) |
| *Angophora* | Smooth-barked Apple<br>(*A. costata*) | 1223 (1002) | 1214 (996) | 237 (197) | 15 (15) |
| *Syncarpia* | Turpentine<br>(*S. glomulifera*) | 1603 (1299) | 1595 (1294) | 314 (255) | 17 (17) |

\* Formerly subgenus *Monocalyptus*
<sup>#</sup> *E. cypellocarpa* samples from both study sites were combined to identify fixed private alleles

Eight koalas at the site were fitted with VHF radiotransmitter collars as part of a monitoring program undertaken by Science for Wildlife, allowing for the regular location of individuals for the purpose of sample collection. We collected scats from five of these koalas on three to six occasions between February and July 2015, while the remaining three were sampled on a single occasion in August 2016. On each occasion, the tree species occupied by the koala was identified and fresh faecal pellets were collected within 4 hours of defaecation from a shade cloth mat (2 m by 3 m) placed on the ground directly beneath the koala and stored at -18 °C.

We collected foliage from 2-5 individuals of each potential food tree species from Mountain Lagoon for DNA extraction and evaluation of candidate barcoding genes. For the DArTseq analysis we required samples from a greater number of individuals, and these were sourced from various locations in the greater Blue Mountains area (Table S1). Leaves were collected from live branches using either a big-shot line launcher or pole pruner. The leaves were stored at -18 C prior to freeze drying.

#### Aireys Inlet, Great Ocean Road, Victoria

In September 2015, 60 koalas were included in a study conducted by the Victorian State Government’s Department of Environment, Land, Water and Planning. The study aimed to assess the fate of koalas translocated from over browsed habitat dominated by *E. viminalis* and *E. obliqua* at Cape Otway to mixed eucalypt forest 90 km away. The study is described in detail elsewhere by Menkhorst and colleagues (2019). We collected faecal pellets from 14 of the translocated koalas on between 3 and 8 occasions over a year post translocation (total n =62). Pellets were either collected opportunistically during koala captures or from plastic mats placed beneath radio-tracked koalas. Pellets were generally frozen at – 18 °C within 2 hours of defaecation and all were frozen within 10 hours.

We identified ten candidate food tree species from the genus *Eucalyptus* in the translocation area (Table 2). We collected leaves from twenty individual trees of each species for DNA analysis using either a big-shot line launcher shot or pole cutter. The leaves were stored at -18 ͦ C prior to freeze drying.

**Table 2:** Species-specific SNPs identified in candidate koala food tree species (genus *Eucalyptus*) at Aireys Inlet.

| Subgenus | Common Name (Species) | Fixed private SNPs (strict) | Subset for Diet Analysis (strict) | Most Suitable for DarTag (strict) | Designed Tags (strict) |
| --- | --- | --- | --- | --- | --- |
| *Eucalyptus* | Messmate (*E. obliqua*) | 1016 (30) | 252 (30) | 60 (6) | 16 (10) |
|  | Brown Stringybark (*E. baxteri*) | 623 (284) | 613 (282) | 144 (75) | 4 (4) |
|  | Narrow-leaved Peppermint (*E. radiata*) | 885 (0) | 68 (0) | 12 (0) | 2 (0) |
|  | Western Peppermint (*E. falciformis*) | 524 (128) | 513 (123) | 123 (21) | 14 (12) |
|  | (*E. radiata* & *E. falciformis*) | 1750 (30) | 294 (30) | 57 (7) | 8 (8) |
| *Symphyomyrtus* | Tasmanian Blue Gum (*E. globulus*) | 727 (23) | 202 (23) | 44 (6) | 10 (4) |
|  | Mountain Grey Gum (*E. cypellocarpa*)# | 696 (9) | 91 (9) | 18 (1) | 10 (2) |
|  | Swamp Gum (*E. ovata*) | 926 (77) | 296 (76) | 61 (16) | 17 (17) |
|  | Scent-bark (*E. aromaphloia*) | 625 (0) | 43 (0) | 7 (0) | 3 (0) |
|  | Manna Gum (*E. viminalis*) | 589 (0) | 5 (0) | 3 (0) | 4 (0) |
|  | (*E. aromaphloia* & *E. viminalis*) | 1153 (6) | 89 (6) | 22 (0) | 3 (0) |
|  | (*E. v. cygnetensis*) | 345 (1) | 46 (1) | 9 (0) | 6 (0) |
|  | (*E.v. viminalis*) | 534 (1) | 14 (1) | 1 (1) | 1 (0) |
|  | Red Ironbark (*E. tricarpa*) | 2675 (1570) | 2618 (1543) | 577 (351) | 16 (16) |

\* Formerly subgenus *Monocalyptus*

### DNA Extractions

DNA was extracted from both the reference leaves and the koala scats using CTAB extraction buffer with chloroform clean-up and ethanol precipitation. Leaf samples were freeze-dried prior to DNA extraction. 25 mg of freeze-dried leaf tissue was ground into a fine powder using two 1.5 mm diameter steel beads in a TissueLyser II (Qiagen) at 30 Hz for 1 minute. 100 mg of each scat sample was also ground into a fine powder using the TissueLyser II (Qiagen) except that each sample was bead beaten three times after snap freezing in liquid nitrogen. The samples were then digested in 500 μl (leaf) or 1000 μl (scats) of extraction buffer (0.1 M Tris-HCL, 1.4 M NaCl, 40 mL of 0.02M EDTA, 20 g.l^-1^ of cetyltrimethyl ammonium bromide and 0.3 % β-mercaptoethanol) in a ThermoMixer (Eppendoff) at 65 °C for 1 hour with mixing at 1000 rpm. The digested samples were centrifuged for 10 minutes at 16 000 g. The supernatant was then washed twice in one volume of chloroform:isoamyl alcohol (24:1). The DNA was then precipitated by addition of a half volume of 5M NaCl to the separated aqueous phase, followed by 3 volumes of cold 95% ethanol and incubation at -20 °C for 1 hour. The samples were then centrifuged for 10 minutes at 16 000 g. The DNA pellet was washed with 700 μl of 70% ethanol followed by 700 μl of 95% ethanol. The DNA was dried for 30 minutes at 65 °C and resuspended in 40 μl of TE buffer. Inhibitors were removed from the DNA extracts using the OneStep PCR Inhibitor Removal Kit (Zymo) according to the manufacturer’s instructions.

### Barcoding genes from candidate food tree species

We assessed the suitability of three nuclear genes (*ITS, ETS* and *CCR*) and two chloroplast genes (*ndh* and *matK*) for food tree species identification in the Blue Mountains koala population (Table 3). A fragment of each gene was individually amplified by polymerase chain reaction (PCR) from the leaf DNA extracts using the MyTaq reaction buffer and polymerase (Bioline) according to the manufacturer’s protocol. The forward and reverse primers (Table 3) were each at a final concentration of 0.2 µM, with 20 ng of template DNA added to a final reaction volume of 20 µl. Amplification was performed with a 3 min denaturation at 95° C, followed by 30 (*ETS*, *CCR*), 35 (*matK*, *ndh*) or 36 (*ITS*) cycles of 30 s at 95°C, annealing for 30 s and a 45 s (*CCR*, *ITS*, *ETS*) or 1 min 30 s (*matK*, *ndh*) extension at 72° C, followed by a final extension of 5 min at 72° C. For the *ITS* gene fragment the annealing temperature for the first 3 cycles was 60° C, followed by 3 cycles at 57° C and the remaining cycles at 54° C. The annealing temperatures for the other genes can be found in Table 3. All genes amplified according to this protocol, except for *ndh.* Attempts to modify the protocol to amplify *ndh* were unsuccessful and thus no analysis of this gene was possible. The PCR products from the other four genes were sequenced on an ABI 3500 Genetic Analyser at the Hawkesbury Institute for the Environment, Western Sydney University.

**Table 3:**
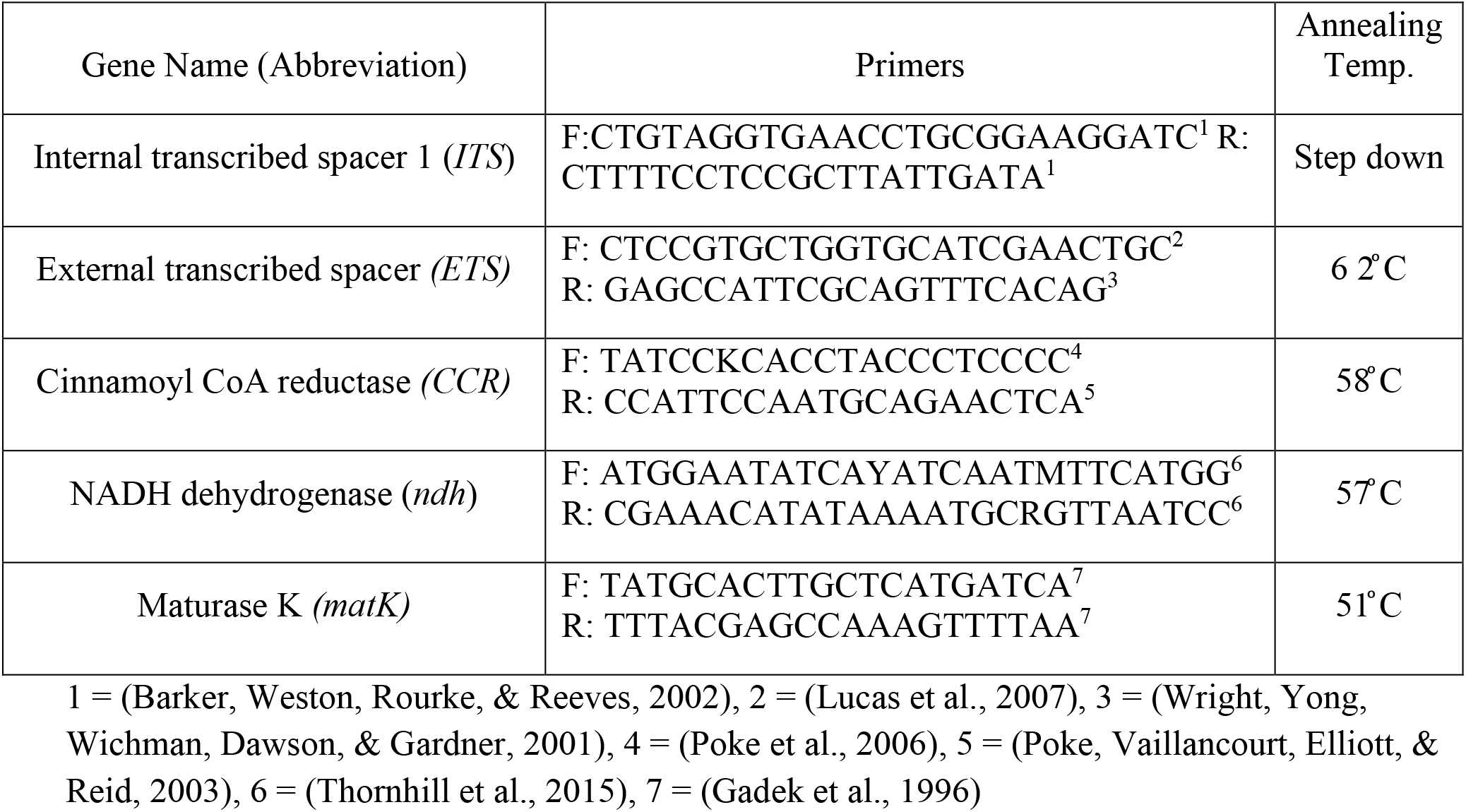
Primers and annealing temperatures for Barcoding genes.

The forward and reverse sequencing tracers were assembled after quality trimming (error probability < 0.05 and fewer than 2 ambiguous nucleotides) in CLC genomic workbench 20.0.01. The assembled sequences were then aligned in CLC (gap open cost = 10, extension cost = 1, end gap cost = free). Short sequences were subsequently manually removed, and the sequences trimmed to a standard length.

The ability of each gene to differentiate the different species of eucalypts was then determined by two measures. Firstly, the polymorphic sites were identified in GenAlEx 6.1 (Peakall & Smouse, 2006, 2012) and manually assessed to determine if any of the alleles were species specific. Secondly, Principal Components Analysis (PCoA) based on the haploid genetic distance matrix was performed in GenAlEx to determine if each species formed a unique cluster.

### Analysis of sampling regimes and data processing protocols for DArTseq

To determine the extent of within-species polymorphism observed in DArTseq data, six trees from each of nine *Eucalyptus* species (Mountain Lagoon: *E. beyeriana, E. cypellocarpa, E. deanei* and *E. punctata;* Aireys Inlet: *E. aromaphloia*, *E. globulus*, *E. obliqua*, *E. ovata* and *E. radiata*) were sequenced at high density (returning approximately 2.5 million reads per sample) on the DArTseq platform by Diversity Arrays Technology P/L, Canberra, Australia (DArT). The *Eucalyptus* Dartseq (1.0) product was selected that uses the PstI/HpaII restriction enzyme combination, which has been optimised for *Eucalyptus* and returns approximately 50 000 loci. Read counts for targets (DNA fragments) and single nucleotide polymorphisms (SNPs) within those targets were determined by DArT using their standard pipelines (Grewe et al., 2015; Kilian et al., 2012). Read proportions for the reference and alternative SNP alleles at each locus were determined using custom code in R studio 1.2.5033 (R-Core-Team, 2012).

To establish if pooling multiple individuals of the same species for sequencing substantially impacted the identification of species-specific SNPs, DNA from the six individuals from each of the nine species described above were pooled by species and re-sequenced on the DArTseq platform.

To assess how increasing the number of individuals sampled per species affected the extent of within-species variation detected, DNA from an additional 13 or 14 trees from each of the nine species were pooled by species and sequenced. Allele calling and comparisons among the pooled and individual samples of each species were calculated using custom written code in R studio 1.2.5033 (R-Core-Team, 2012).

### Identification of species-specific SNPs using DArTseq

To identify species-specific SNPs (i.e. those only found in the target species) for the twelve potential food tree species at Mountain Lagoon and 10 species at Aireys Inlet, DNA from 17-20 trees from each of the remaining 13 species not sequenced above were pooled by species and sequenced on the DArTseq platform.

Prior to identification of species-specific SNPs, a Principal Components Analysis was performed on the Hamming genetic distance matrix generated from the SNP data in GenAlEx 6.1 to confirm the species designations of the samples. Heterozygous loci were then removed from the read count table as these loci could not contain fixed private alleles. Species-specific SNPs were identified in the homozygous loci using custom written code in R studio 1.2.5033 (R-Core-Team, 2012).

### Diet determination from species-specific markers

#### DArTseq

To trial if DArTseq was a viable platform for diet determination, six scats sampled from Mountain Lagoon koalas were processed using Eucalyptus DArTseq (1.0) product and sequenced at high density (returning approximately 2.5 million reads per sample). All reads that corresponded to the species-specific SNPs identified from the analysis of the eucalypts were then extracted from the sequencing data by DArT using a custom bioinformatics pipeline (two independent algorithms written in R v3.5.0. and Perl v5.26.0 returning the same results). Those reads were then tallied by species to estimate diet composition.

#### DArTag

The list of species-specific SNPs identified from the DArTseq analysis of the potential food tree species from Mountain Lagoon and Aireys Inlet were filtered to retain only a single SNP per unique DArT sequencing fragment in order to minimise the impact of linkage disequilibrium among potential markers. Those SNPs that had < 95% reproducibility or were detected in less than 50% of individuals from the target species were also excluded. A total of 312 SNPs (2 to 67 per species) were then selected for oligo design. Oligos were designed for the selected SNPs in house by DArT by identifying probe regions flanking the selected SNPs using primer design software in Genious (Drummond et al., 2009). Of the candidate species-specific SNPs only those that were found in the middle of the DArT target were suitable for oligo design. Of the suitable SNPs, those that were fixed, had higher DArTseq read counts and higher reproducibility were preferentially selected. Where SNPs from DArT targets that were only present in some individual trees had to be included, those that were present in a higher proportion of the individuals were given preference.

The selected tags (SNPs and flanking regions) were then amplified and sequenced from four individual trees of each species using the DArTag platform to establish that they remained species specific on the different platform. Briefly, the pooled species-specific oligos were hybridized to denatured eucalyptus DNA, then the targeted SNPs were copied and amplified with simultaneous addition of demultiplexing primers. The products of DArTag assay were then sequenced on the HiSeq 2500 (Illumina), demultiplexed and targeted SNPs detected using DArT P/L’s proprietary analytical pipeline (DArToolbox). Those SNPs that were confirmed to be species-specific on the DArTag platform were amplified and sequenced from the faecal DNA extracts (as described above) in order to determine the composition of the koalas’ diets. When only a single read was returned for a species it was pruned from the resulting dataset.

## Results

### Molecular barcoding genes for potential dietary eucalypts

None of the four candidate barcoding genes tested were able to successfully delineate the twelve potential food species present at Mountain Lagoon (Fig. 1). However, all four genes did separate the different genera and subgenera of *Eucalyptus*. In the PCoAs of the four genes, the two subgenera of *Eucalyptus* were found to be distinct and the ironbarks clustered separately from the other members of subgenus *Symphyomrtus*.

**Fig 1:**
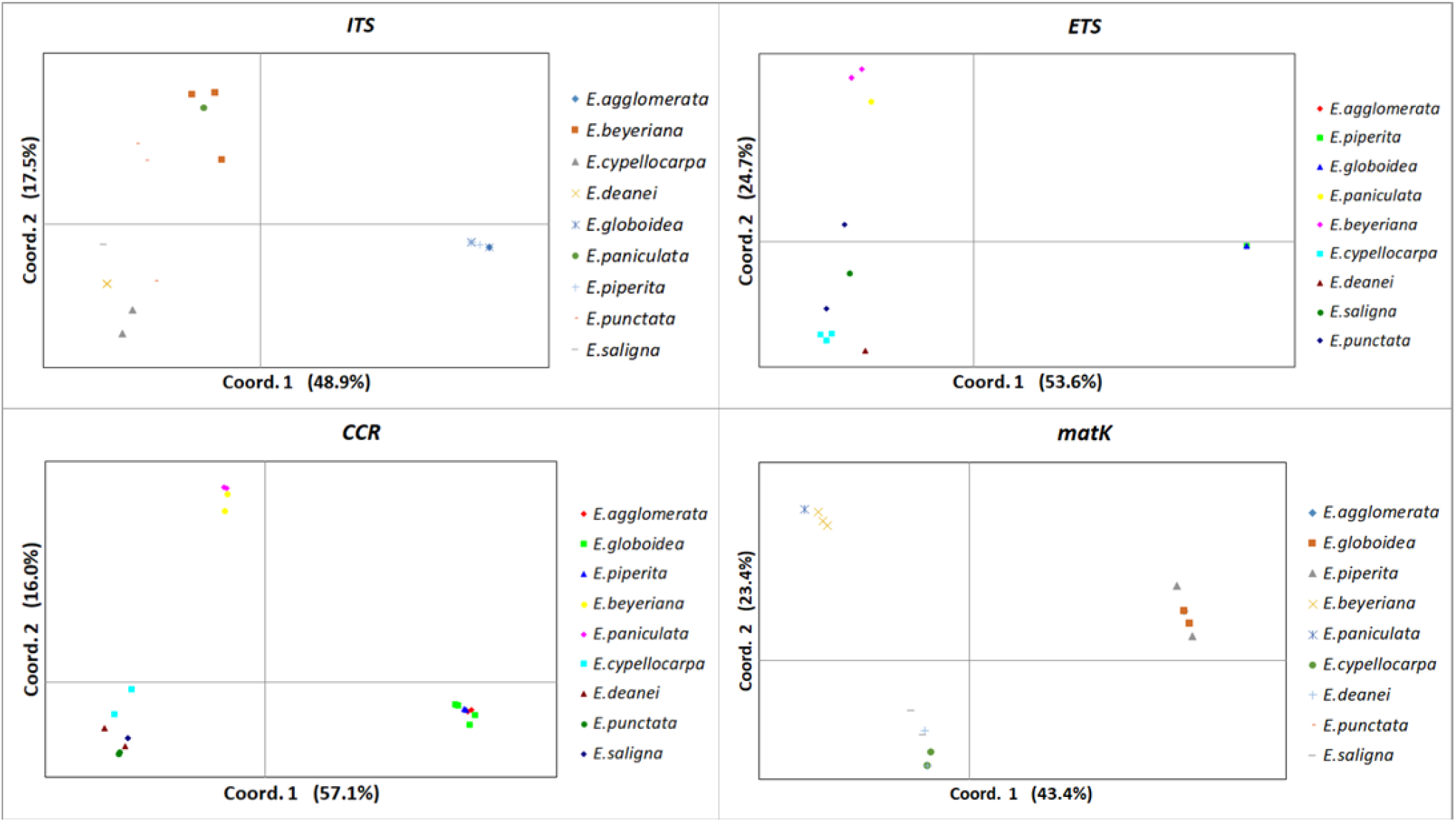
The first two axes of the principal components analyses based on the haploid distance matrices of each of the four candidate barcoding genes for the nine candidate koala food trees from the genus *Eucalyptus* at Mountain Lagoon.

After quality control, 33 polymorphic sites were identified in the 288 nucleotide *ITS* sequence. Of those none were private to any of the *Eucalyptus* species, though six fixed private SNPs were identified for *S. glomulifera*, one for *A. costata* and two for *C. gummifera*. Additionally, one SNP was specific to the subgenus *Eucalyptus* and one separated the ironbarks from the other *Symphyomyrtus* species.

Over the 338 bp length of the trimmed *ETS* sequence 86 polymorphic sites were identified. As for *ITS* none were private to any of the *Eucalyptus* species, though 30 fixed private SNPs were identified for *S. glomulifera* and 23 for *A. costata*. No *ETS* sequences for *C. gummifera* were retained for analysis after quality control. Five SNPs were specific to the subgenus *Eucalyptus* and three separated the ironbarks from the other *Symphyomyrtus* species.

The *CCR* gene could only be amplified in the *Eucalyptus* species. Fifty-six polymorphic sites were identified across the 429 bp sequence. The different potential dietary species could be better resolved with the *CCR* gene than the other genes tested, with one fixed private SNP identified for *E. paniculata*, two for *E. saligna* and one for *E.cypellocarpa.* However, the remaining six species could not be resolved. Fifteen SNPs were specific to the subgenus *Eucalyptus,* seven were specific to the ironbarks and three were specific to the other *Symphyomyrtus* species.

As expected for a chloroplast gene, *matK* showed considerably less sequence variation than the other genes, with only 38 polymorphic sites identified over the 839 bp sequence. None of the *Eucalyptus* species could be resolved, although *A. costata* was separated from the other species by five SNPs and a six bp indel. *MatK* sequences were not obtained for *S. glomulifera* and *C. gummifera* as the gene showed little promise for use in diet determination. Six SNPs were specific to the subgenera *Eucalyptus* and three separated the ironbarks from the other *Symphyomyrtus* species.

### Analysis of sampling regimes and data processing protocols for DArTseq

A total of 321 498 validated SNPs were identified from the 22 candidate food tree species at Mountain Lagoon and Aireys Inlet. DArTseq data produces four possible genotypes: 0) the locus is not detected; 1) the reference allele is detected; 2) the alternative allele is detected; or 3) both alleles are detected. Across all the detected loci in the individual trees (excluding the pooled samples), in 66.2% of cases only the reference allele was detected, while in 32.5% of cases only the alternative allele was detected. In the remaining 1.3% of cases both alleles were detected. The proportion of reads assigned to the alternative allele, when both alleles were detected, appeared to follow an approximately normal distribution centred on 50% (Fig. S1), as is expected in the case of heterozygous loci in a diploid organism. However, the full range of proportions were observed (0 < proportion of reads alternative allele < 1) and there was no clear delineation between a homozygous and heterozygous locus based on read proportions (Fig. S1). Therefore, given the small proportion of cases (loci x individuals) where both alleles were detected, all such instances were scored as heterozygous. By this definition 45.5% (146 461) of loci were heterozygous in at least one individual tree.

When the genotypes of six individual trees from each of nine species were compared to the pooled genotypes of those samples (the same six individuals from each species pooled by species prior to sequencing) then the same species level genotype was found in 53.5% of cases across all loci (i.e. if the individual trees were all homozygous for an allele then the pooled sample was also homozygous for that allele; or if both alleles were detected among the six trees then the pooled sample was heterozygous; Fig. S2). The 46.5% of instances where the individual samples did not match the pooled samples could be separated into two different types of mismatches: 1) the locus was only detected in some individuals from the species, and this could not be detected in the pooled sample; or 2) the set of alleles found in the individual samples did not match those found in the pooled sample (e.g. both the reference allele and the alternative allele were detected among the six individual trees but the pooled sample was homozygous). Type 2 mismatches were rare in the dataset and found in only 1.9% of comparisons. It was observed that the frequency of type 1 mismatches decreased as the average number of reads for the locus increased (Fig. S2), which suggested that the absence of a locus in an individual may be due to locus dropout and not a true absence from the individual. The estimated frequency of type 1 mismatches attributable to true absences was taken as the type 1 mismatch frequency for loci with > 200 reads, 9.0%, as it was considered that the dropout rate for these loci was likely to be very low. Using this estimate the dropout rate was calculated to be 7.5% per locus per individual across all loci, producing an estimated type 1 mismatch frequency due to dropouts of 35.2% across the dataset (Fig. S2).

On average across all loci in the nine test species the pooled sample of six individuals had the same genotype as the pooled sample of fourteen trees in 85.1% of comparisons (range by species: 81.9% to 91.0%). In 12.4% of cases the locus was not detected in the pool of six or the pool of fourteen individual trees when an allele was detected in the other sample. In the remaining 2.5% of cases (range by species: 0.9% to 3.7%) the pools of six and fourteen had different genotypes. In 60.5% of those instances (or 1.5% of total comparisons) the pool of fourteen trees revealed an allele that was not detected in the pool of six individuals.

### Identification of species-specific SNPs using DArTseq

The principal components analysis of the Hamming genetic distances for the candidate food tree species revealed that the three non-*Eucalyptus* species were distinct based on their multi-locus SNP genotypes and that the two subgenera of *Eucalyptus* formed distinct clusters (Fig. S3). Additionally, the ironbarks formed a separate cluster to the other *Symphyomyrtus* species. When a separate PCoA was performed on each of the macro-clusters, it was evident that the different species were generally well separated (Fig. 2). However, among the species that belonged to subgenus *Eucalyptus, E. radiata* and *E. falciformis* clustered together as did *E. agglomerata* and *E. globoidea*. Among the non-ironbark *Symphyomytus* species *E. deanei, E. punctata* and *E. saligna* were clearly separated from the other species (Fig. 2). Once those three species were removed from the PCoA then the separation among the remaining species could be observed. However, *E. aromaphloia* and *E. viminalis* showed some overlap (Fig. 2).

**Fig 2:**
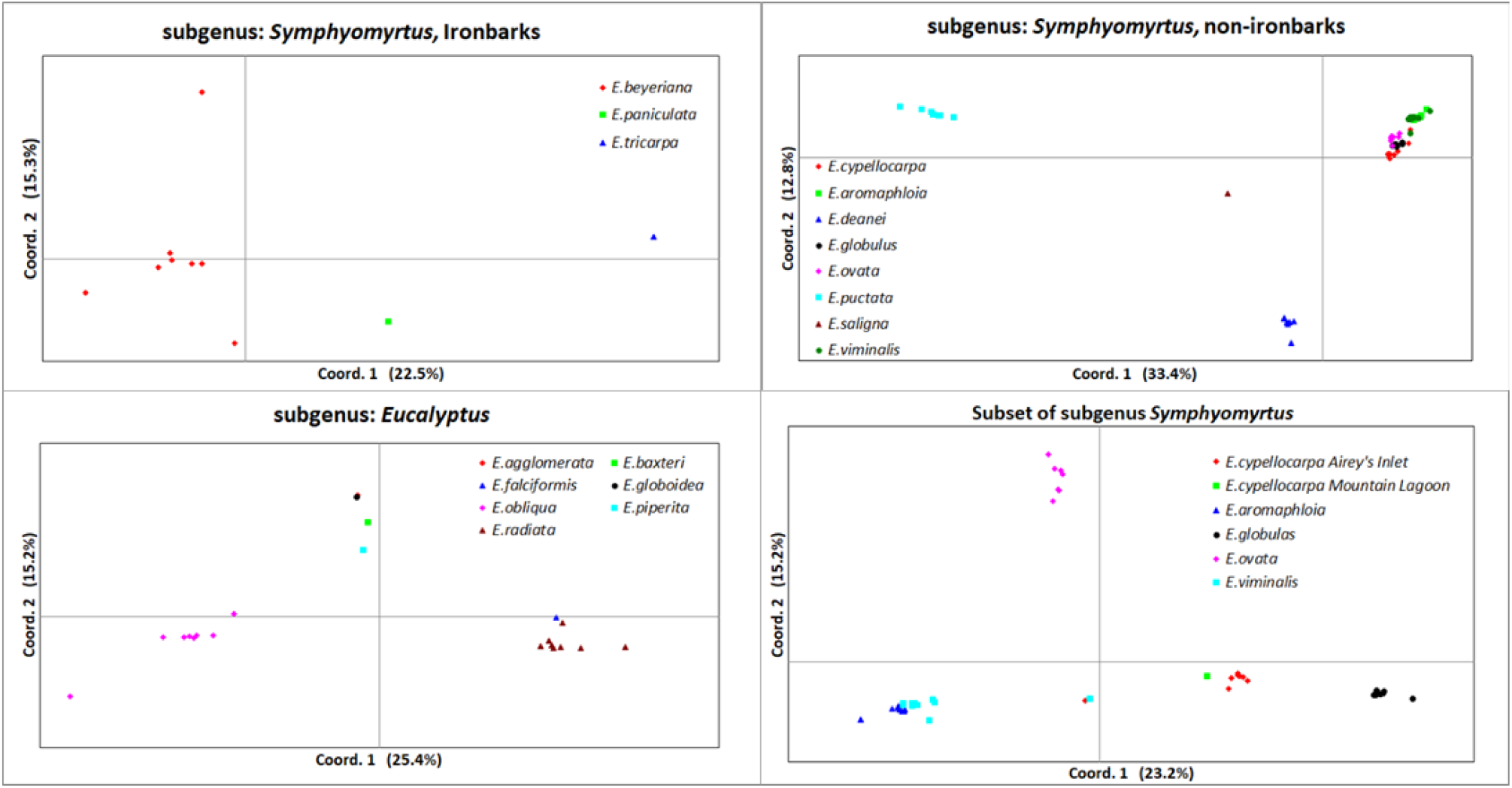
The first two axes of the principal components analyses based on the Hamming distance matrix generated from DArTseq SNPs data for subsets of the candidate koala food trees from Mountain Lagoon and Aireys Inlet.

The *E. cypellocarpa* samples from Aireys Inlet clustered with the *E. cypellocarpa* samples from Mountain Lagoon in the PCoA (Fig. 2). Therefore, these two populations were pooled for the analysis to identify the species-specific SNPs. Two individual tree samples, one labelled as *E. viminalis* and one labelled as *E. cypellocarpa,* did not cluster with their respective species but instead clustered together mid-way between the two species clusters (Fig. 2). We considered that these samples were potentially *E. viminalis/E. cypellocarpa* hybrids and they were excluded from the analysis to identify the species-specific SNPs.

A total of 29 601 fixed private alleles were identified across the 22 potential dietary tree species from the two study sites (Table S2). Of those, an absence of the locus was fixed in 1 229 cases. As the absence of a locus cannot be used as a reliable species-specific marker for detecting diet composition from scat samples, those alleles were excluded from further analysis.

As a high rate of allele dropout was estimated for the dataset (see above), two types of private SNPs were considered. A ‘strict’ species-specific SNP was one that was detected in all individuals of the target species and not detected in any other species. A ‘null allowed’ species-specific SNP was one where the locus was not detected in all individuals of the target species but when it was detected all individuals of the target species had that SNP. Following these definitions, 8 818 strict SNPs and 19 554 null allowed SNPs were identified across the species.

The number of SNPs identified for each potential food species varied from 345 to 2746 (Tables 1 and 2) and no strict SNPs were identified for *E. aromaphloia*, *E. viminalis* or *E. radiata*. In an attempt to improve our ability to detect these species in the scats, we identified SNPs that were private to *E. radiata* and *E. falciformis,* as well as SNPs that were private to *E. viminalis* and *E. aromaphloia* (Table 2). Additionally, during sample collection it was observed that two subspecies of *E. viminalis* (*viminalis* and *cygnetensis*) were present at Aireys Inlet and private SNPs were identified for each of these subspecies (Table 2).

After removal of linked SNPs and those with low reproducibility a total of 15 604 (52.7%) were retained for diet determination using DArTseq and for DArTag design, although the proportion of SNPs retained varied greatly among species (Tables 1 and 2). Of those, 6102 had a suitable SNP position (adequate flanking sequence) for DArTag design and of those approximately 3321 had the more ideal primer design sites based on the primer design pipeline (Tables 1 and 2).

### Diet determination from species-specific markers

#### DAr*Tag*

On examination of the initial DArTag data returned from the reference tree species (four individuals per species), it was revealed that only 45.3% of the selected fixed private SNPs identified in the DArTseq analysis successfully amplified and were private on the DArTag platform. This relatively low conversion rate can likely be attributed to methodological differences between the platforms, in particular the presence of a restriction enzyme digestion step in DArTseq that is absent from the DArTag protocol. Therefore, the DArTag data were independently analysed without reference to the originally designed markers to identify private SNPs. This analysis identified 58 informative SNPs for the tree species present at Aireys Inlet and 182 for those from Mountain Lagoon. This included SNPs that were private to several combinations of the different species (Table 4) and were included to improve our ability to detect the presence of those species. The compositions of the koalas’ diets were extrapolated from these group SNPs following a set of logical rules that also took into account the reliability of the markers for each group (online supplement 1).

**Table 4:**
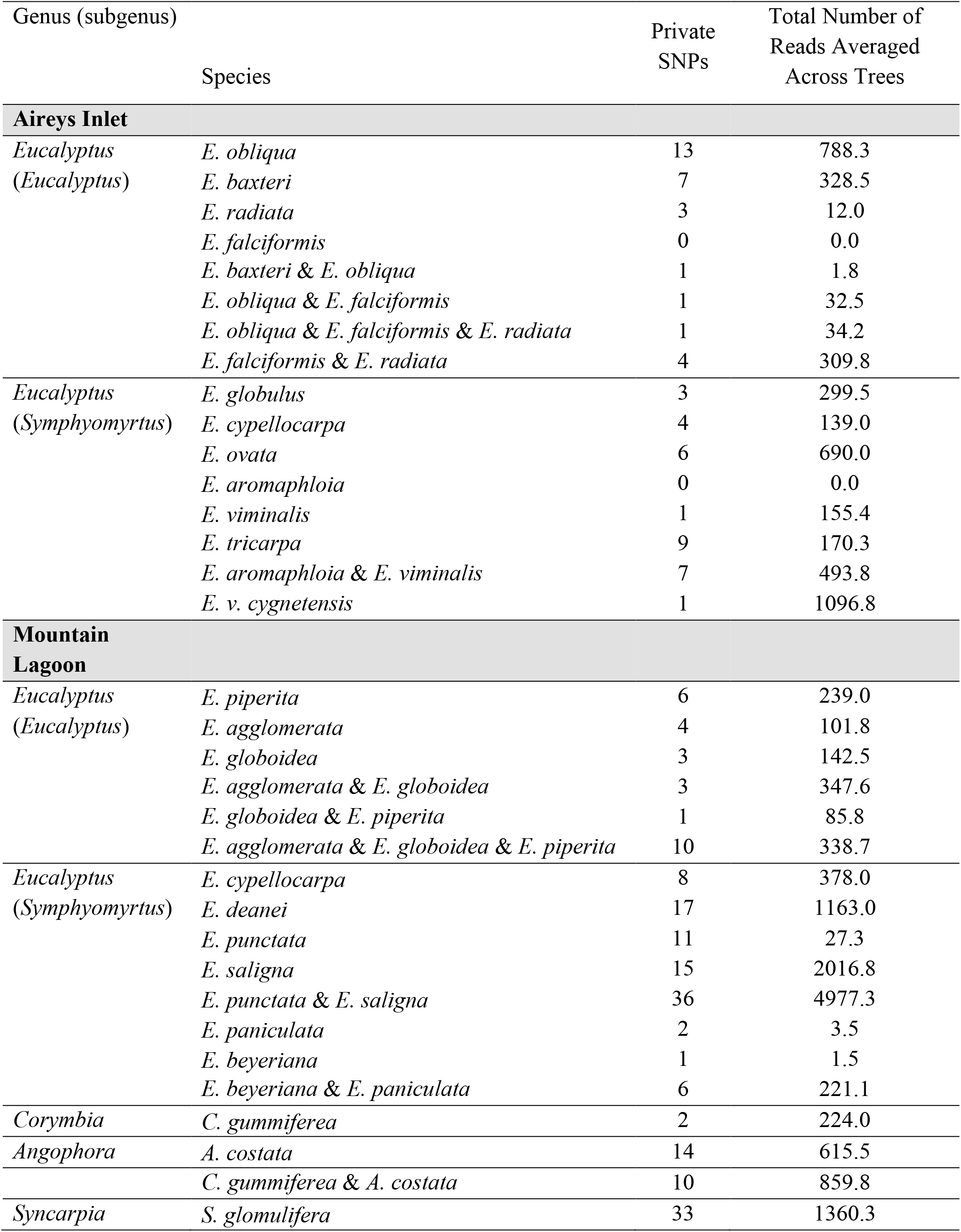
Numbers of species-specific SNPs and reads from leaf DNA from the DArTag analysis for the candidate koala food tree species.

For the majority of species, we identified several private SNPs that amplified well (Table 4). However, we were only able to identify a small number of private SNPs for *E. aromapholia, E. viminalis, E. radiata* and *E. falciformis* at Aireys Inlet, such that our ability to detect these species was relatively poor. Nevertheless, we had a strong panel of SNPs to detect *E. aromapholia* and/or *E. viminalis* as well as *E. radiata* and/or *E. falciformis.* Similarly, at Mountain Lagoon our ability to detect *E. paniculata* and *E. beyeriana* was relatively poor, nonetheless, we had six SNPs that amplified well to detect the presence of ironbarks (*E. paniculata* and/or *E. beyeriana*) in the diet (Table 4). Although we identified 11 SNPs for *E. punctata,* these amplified relatively poorly. We did, however, assemble a strong panel of SNPs to detect the presence of *E. saligna* and also to detect *E. punctata* and/or *E. saligna.* Therefore, in most instances, the presence of *E. punctata* in a sample could be inferred from the presence of the combined SNPs and the absence of *E. saligna* specific SNPs.

The number of reads returned from the target tree species varied between loci (Table 4), in part due to differences in the efficiency of hybridization and amplification prior to DArTag sequencing. The number of reads returned from the target tree species for each SNP was moderately correlated with the number of reads returned from the scat samples in which that SNP was detected (Fig. 3; R^2^ = 0.52). Therefore, in order to estimate the relative proportions of the different tree species in the koalas’ diets, the raw DArTag reads from the faecal pellets were scaled by the total number of reads returned across the private SNPs in the trees (averaged across all individuals from the target species).

**Fig. 3:**
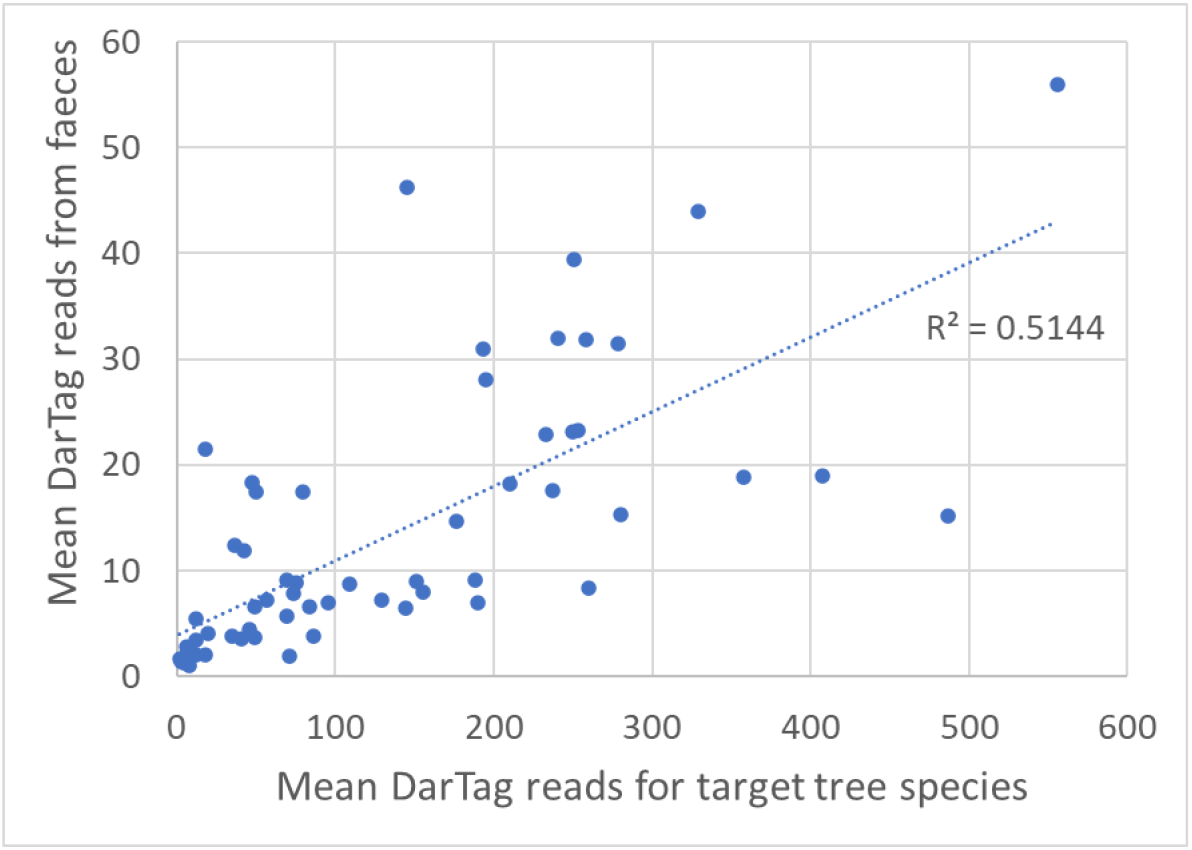
Correlation between the average number of DArTag reads returned for leaf DNA from the target trees species and the average number of reads detected in faecal samples for private SNPs detected in ≥ 5 samples.

#### Comparison of DArTseq and DArTag

An average of 20.2 reads (range: 5 – 39) containing species-specific SNPs where recovered from the six faecal DNA samples when they were run on the DArTseq platform, from among the 2.5 million total reads returned per sample on average. By comparison an average of 432 reads (range: 244-979) containing species-specific SNPs were recovered from the same six faecal samples when run on the DArTag platform, from among 50,000 reads on average per sample.

Between two and five species were detected in each sample by DArTseq, while two to seven were detected by DArTag. There were seven instances out of 26, where a tree species was identified in a sample on the DArTag platform that was not detected with DArTseq. In general, those species that were not detected in the DArTseq analysis were estimated from DArTag to be a minor proportion of plant DNA in the pellet in question (average = 7.2%; range = 0.8% – 25.2%). There were two instances where a species was detected by DArTseq that was not detected with DArTag. However, in both cases only a single DArTseq read was identified for the species and thus it was likely to be sequencing error.

There was generally good agreement between DArTseq and DArTag in the estimated proportions that each species represented in the koalas’ diets (R^2^ = 0.80; Fig. 4). However, DArTseq appeared to generally underestimate or fail to detect the minor components of the diet, biasing compositional estimates towards the major components, most likely due to the very low species-specific read counts returned by DArTseq.

**Fig 4:**
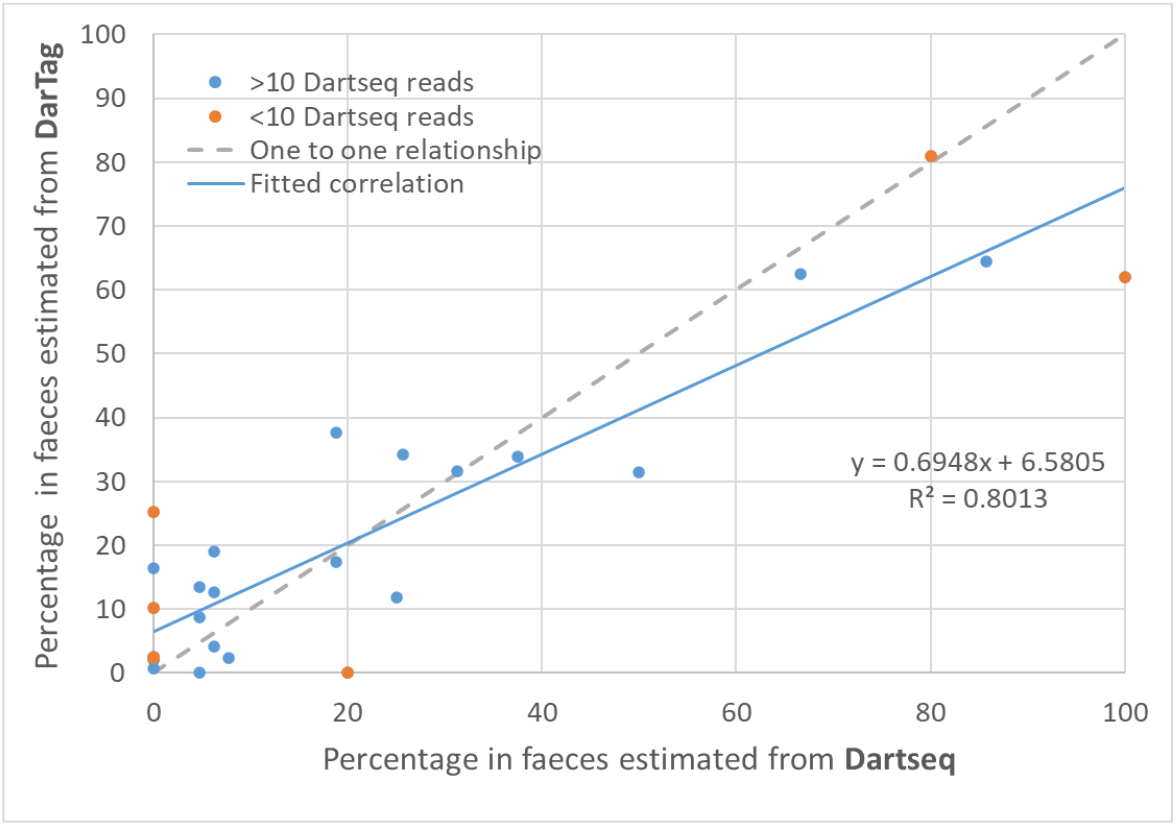
Relationship between proportional representation of tree species in faecal samples as estimated from DArTseq and DArTag. Each point represents a species in a sample.

#### Diets of captive koalas with known diets

Only SNPs for *E. viminalis/E. aromaphloia* were detected from the three captive koalas that were exclusively fed *E. viminalis.* Only *E. obliqua* SNPs were detected from one of the koalas that was fed *E. obliqua*. However, *E. baxteri* was estimated to account for 1.2% and 1.6% of the diets for the other two koalas nominally fed only *E. obliqua. E. baxteri* is another stringybarked eucalypt present at Cape Otway, which can easily be mistaken for *E. obliqua* and presumably accidently fed to those koalas. Two reads containing *E. radiata* SNPs were detected from one of the *E. obliqua* koalas. Because *E. radiata* could not have been mistakenly fed, these reads were likely to be sequencing errors or low-level contamination. However, as the *E. radiata* SNPs amplified poorly these reads were up scaled resulting in *E. radiata* being estimated to form 23.1% of the diet, despite accounting for only 0.4% of SNP reads in the samples.

#### Diet of Mountain Lagoon koalas

Across the eight koalas sampled at Mountain Lagoon, those species that were identified in the scats of a greater number of koalas, and in a higher proportion of pellets, also accounted for a greater average percentage within pellets (Fig. 5a). Although, the proportion of pellets and koalas in which a species was found generally overrepresented rare species in the diet compared to the average percentage within pellets.

**Fig 5.**
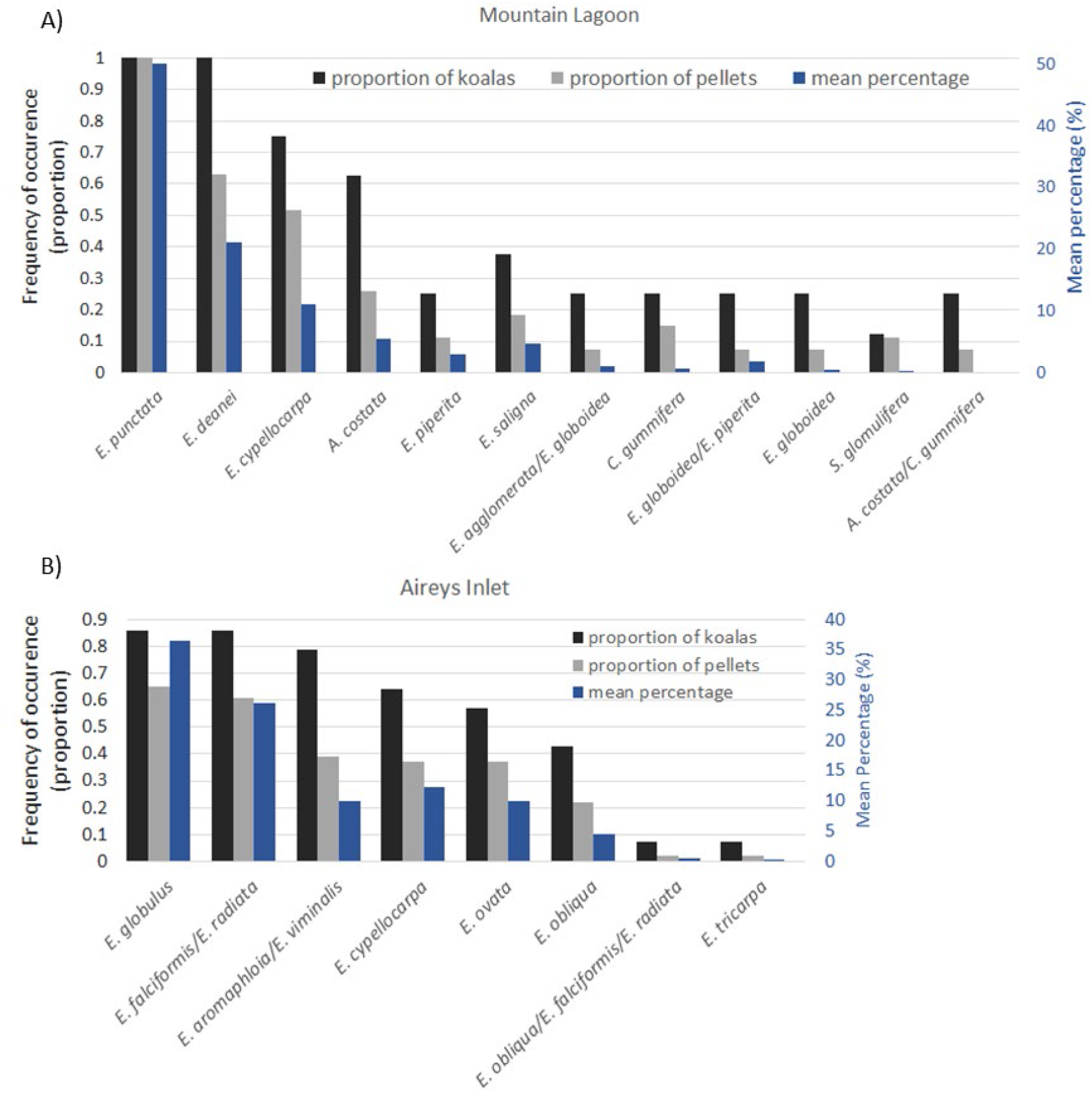
The proportion of koalas that each tree species was detected in and the mean percentage of the diet that they constituted as estimate from DarTag for (A) Mountain Lagoon and (B) Aireys Inlet (post-translocation).

*Eucalyptus punctata* was detected in all samples from all eight koalas and was the dominant component (> 50%) of samples from six of the koalas on at least one occasion (Fig. 6). *Eucalyptus deanei* was also detected in all koalas and dominated or was a major component (> 20%) of at least one faecal sample from five koalas. *E. cypellocarpa* was detected in six koalas and was a dominant or major component of at least one faecal sample from three koalas*. Eucalyptus piperita* and *A. costata* were major components of faecal samples from two and one koalas, respectively. *Eucalyptus saligna, C. gummifera* and *E. globoidea* were detected as smaller components of samples from three, two and two koalas respectively. The non-eucalypt *Syncarpia glomulifera* (turpentine) was detected at very low levels in the faeces of a single koala, on three separate occasions.

**Fig. 6:**
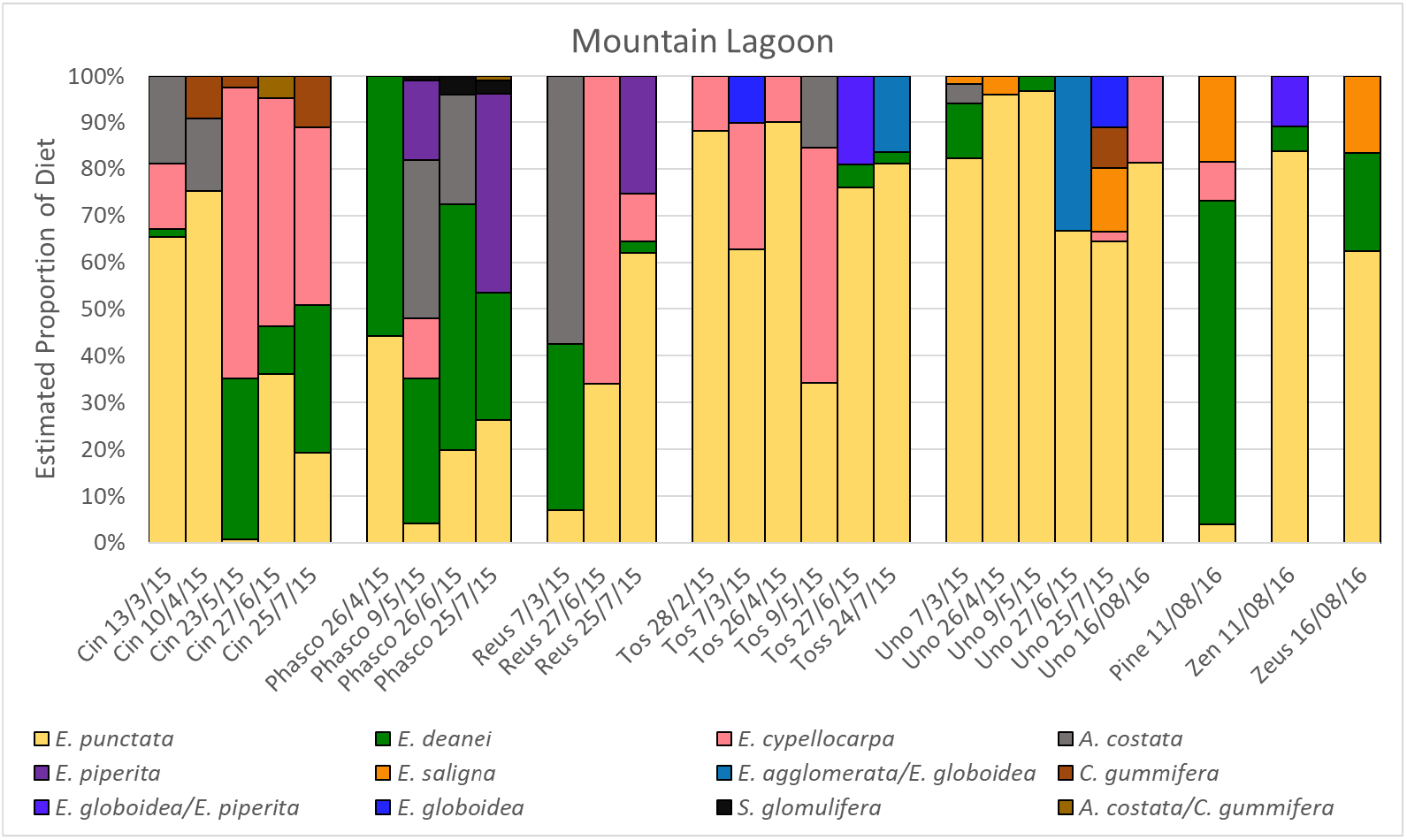
Diet composition for koalas from Mountain Lagoon as estimated using the DArTag platform.

*E. agglomerata* SNPs were not detected in any of the samples. *E. agglomerata/E. globoidea* SNPs were detected in the absence of either *E. globoidea* or *E. agglomerata* SNPs. However, these likely indicate the presence of *E. globoidea* rather than *E. agglomerata* in the diet as they were detected in samples from koalas that were found to be feeding on *E. globoidea* at other time points (Fig. 6). There was no evidence that any of the koalas were feeding on *E. beyeriana* or *E. paniculata,* which were uncommon and unevenly distributed at the site.

Five koalas were sampled on multiple occasions. Samples from three of these koalas were consistently dominated by a single food tree species (either *E. punctata* or *E. deanei*), while the composition of samples from the other two koalas was more variable.

#### Diet of Aireys Inlet koalas

Of the 62 samples from the translocated koalas, six were removed from the analysis as they returned fewer than 5 reads containing private SNPs from the DarTag sequencing. This included four samples collected pre-translocation. Of the remaining pre-translocation samples, collected from ten koalas, eight were dominated (> 60 %) by *E. viminalis/E. aromaphloia* (Fig. 7). While *E. viminalis* and *E. aromaphloia* could not be distinguished, *E. aromaphloia* was not present at the pre-translocation site. *E. obliqua* was also detected in pre-translocation sampled from five koalas and dominated two of them. One pre-translocation sample also contained *E. globulus* and *E. cypellocarpa* SNPs.

**Fig. 7:**
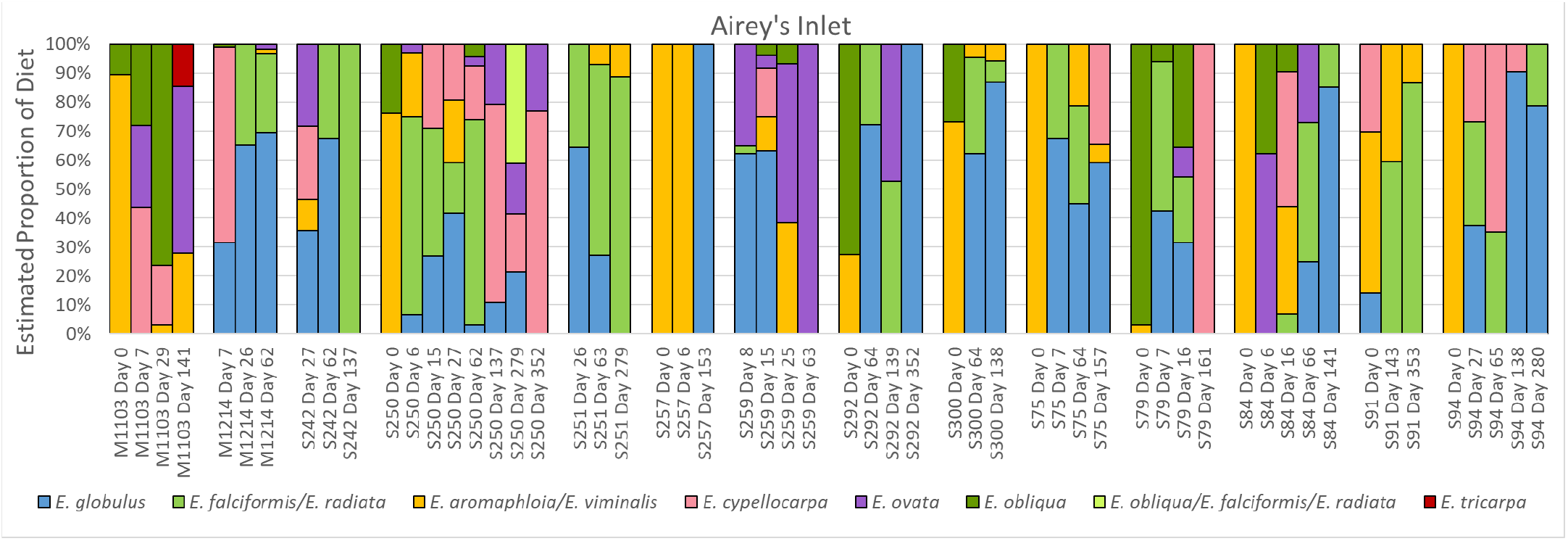
Diet composition for koalas from Aireys Inlet as estimated using the DArTag platform.

After translocation to Aireys Inlet, the koala diets became more species-rich and variable among individuals (Fig. 7). *E. aromaphloia/E. viminalis* no longer dominated the diet of these koalas, although they were still commonly eaten and formed a major component (>20%) of some samples for seven koalas. Both *E. viminalis* and *E. aromaphloia* were available and occupied by koalas at the post-translocation site, so it is likely that both species were eaten.

Similar to Mountain Lagoon, those species that were identified in the scats of a greater number of koalas post-translocation, and in a higher proportion of pellets, also accounted for a greater average proportion within pellets (Fig. 5b). *E. globulus* was dominant in at least one faecal sample from each of ten koalas, while *E. cypellocarpa* was dominant in at least one sample from four of the fourteen koalas. Based on our understanding of the distribution of the tree species and the movement patterns of these koalas, the six samples (from four koalas), that were dominated by *E. radiata/E. falciformis* are likely to indicate feeding on *E. radiata*, a known koala food tree species (Martin, 1985). *E. ovata* and *E. obliqua* were also commonly found as major components of koala faecal samples (Fig. 7). Two reads containing *E. tricarpa* SNPs were detected in a single sample, and *E. baxteri* SNPs were not detected on any occasion.

## Discussion

### Barcoding genes from candidate food tree species

Molecular barcoding genes have been successfully used to characterise the diets of many herbivores and predators from faecal samples (Castle et al., 2020; Goldberg et al., 2020; Kartzinel et al., 2015; King & Schoenecker, 2019; Srivathsan et al., 2016). However, we have shown that such genes are unable to resolve the components of koala diets to species level. This is unsurprising, as these genes have only been successfully deployed by systematists (e.g. Gibbs, Udovicic, Drinnan, & Ladiges, 2009; Steane, Byrne, Vaillancourt, & Potts, 1998) to investigate higher-level phylogenetic relationships among eucalypts, and whether their sequences are fixed or private has not previously been assessed by sampling multiple individuals per species. The koala is unusual in being an obligatory specialist mammalian herbivore (Shipley et al., 2009) which feeds on numerous congeneric species throughout its geographic range, and these co-exist alongside hundreds of other congeneric species that are not consumed. Therefore, species level identification of dietary components is critical for the assessment of suitable koala habitat. Additionally, as the nutritional quality and chemical defences of eucalypts, including koala food trees, varies substantially among genera, subgenera and species, high taxonomic resolution becomes important if diet composition analysis is to provide insight into koala nutrition (DeGabriel, Wallis, Moore, & Foley, 2008; Marsh et al., 2019; Marsh et al., 2020; Moore, Foley, Wallis, Cowling, & Handasyde, 2005). As such, molecular barcoding genes appear to have limited utility in this system, except perhaps in regions where potential dietary species belong to different subgenera or genera. Yet, even in such circumstances it should be considered that barcoding gene markers are typically much longer (>250 bp) than SNP based markers (∼50 bp) and may be harder to amplify from faecal samples where the dietary DNA is likely degraded.

### Determination of suitable sampling and data processing protocols for DArTseq

This is the first application of DarTseq to diet analysis from faecal samples and before any new molecular method can be utilised it is important to ascertain suitable sampling and analysis protocols to ensure robust and reliable conclusions can be obtained. Our assessment of the DArTseq platform for the identification of species-specific SNPs revealed that the heterozygosity rate within eucalypt species was very low. Additionally, increasing the sample size from six to twenty individuals per species only revealed additional within-species diversity in 1.5% of cases. From these findings it can be concluded that it is only necessary to sample a small number of individual trees of each species at a site in order to capture the majority of the genetic diversity present. This may be because DArTseq preferentially identifies SNPs in functional regions of the genome that are frequently subject to purifying selection (Heller-Uszynska et al., 2011), reducing within-species diversity compared to other genetic markers such as microsatellites that are found in non-coding regions.

While the low within-species diversity present in DArTseq SNP markers suggests that pooling multiple individuals of a species is not necessary to increase sample sizes, such pooling does not substantially decrease data quality. The actual mismatches between the genotypes returned from pooling six individuals and sequencing those individuals separately was very low. This suggests that rare alleles are not generally missed when pooled samples are used.

Although low species diversity does not necessitate the sequencing of multiple individuals of a species, the high allele dropout rate observed in the DArTseq data (estimated at 9%) makes such sampling critical. The high dropout rates present in the DArTseq data suggest that the absence of a locus is not a reliable species-specific marker. However, such markers cannot be used when identifying dietary species from faeces and thus such dropout rates pose less of an issue for this application than might be the case for other analyses. Nonetheless, sampling multiple individuals from a species will reduce the effect of dropouts on species level diversity. Further, it is important to take the high dropout rate into account during analysis. In this case we did so by considering ‘Null Allowed” SNPs when identifying potential species-specific markers.

The detection of two erroneous *E. radiata* SNPs in one of the captive koalas with known diet leads us to suggest that further steps should be taken to limit the effect of noise in the dataset on the study’s conclusions. We note that among the samples taken from Aireys Inlet koalas, *E. radiata* SNPs were only detected where *E. radiata/E. falciformis* SNPs were also found. Therefore, one approach would be to only consider a species as present in a sample if more than one SNP for that species is detected. Further, removal of low frequencies reads (e.g. < 5) would be advisable and is commonly used in other NGS applications. While such filtering of the dataset will reduce the detection of low frequency components of the diet, it will improve confidence in the findings, and it can be argued that the rare components of the diet do not play a major role in host nutrition and feeding ecology.

### Identification of species-specific SNPs using DArTseq

In general, the DArTseq platform returned a large number of species-specific SNPs and, unlike the molecular barcoding genes, provided species-level resolution of candidate koala food trees. However, there were several pairs of species that could not be separated on the basis of private alleles and had overlapping genotypes. This was particularly true for several of the species at Aireys inlet, where the number of SNPs per species was generally lower than for those at Mountain Lagoon. This difficulty in identifying suitable private alleles is likely due to incomplete linage sorting among closely related species and recent hybridisation, which are both common among eucalypts (Jones et al., 2016; Steane et al., 1998). The reduced resolution seen at Aireys inlet may be due to more frequent hybridisation among species at that site. Indeed, we identified two putative hybrid individuals among the samples we collected for this study. Nonetheless, the DArTseq method was superior to the use of barcoding genes in this system.

### Diet determination from species-specific markers

By identifying species-specific SNPs in faecal samples, we were able to successfully characterise the diets of koalas at both our study sites. In general, our genetic characterisation of the diets of captive koalas were consistent with the species fed, in support of the validity of this approach. Additionally, the species that were identified as major and minor components of the koalas’ diets at both sites were in broad agreement with previously identified preferred koala food tree species. For instance, *E. punctata, E. deanei* and *E. cypellocarpa* are listed as high use trees in the Blue Mountains region (OEH, 2018) as are *E. globulus* and *E. cypellocarpa* in southern Victoria (Martin & Handasyde, 1999; Mitchell, 2015). Diet composition estimates from our method for the Mountain Lagoon koalas can be compared to patterns of tree use by the same koalas as reported by Gallahar et al. (2021). For most koalas, the species most commonly used by koalas during the day were also dominant food species (especially *E. punctata*). An exception was *C. gummifera,* which was heavily used by koala *Cin* but consumed sparingly. However, several species accounted for more of the diets than tree use patterns suggested. This was especially true of *E. cypellocarpa* (e.g. *Tos, Reus*). The diet of koala *Phasco* was also more species-rich than suggested by day-time tree use observations.

Due to the significant differences among food tree species in nutritional quality, quantitative estimates of diet composition are essential to enable nutritional inference for koalas. All diet composition methods that rely on postingestive samples are subject to quantitative error as a consequence of the differential digestibility of different diet items (Garnick, Barboza, & Walker, 2018). DNA barcoding techniques have been suggested to be particularly susceptible to this problem, with quantification also complicated by the potential for differences in DNA density between tissues, and varying barcoding gene copy numbers (Pompanon et al., 2012). These difficulties are lessened by the koala’s specialised diet, as different diet items (foliage of different Myrtaceous tree species) can be expected to differ less in these respects than, for example, grasses vs browse, or animal vs plant tissue. Additionally, the SNP markers identified in this study were located in the nuclear genome and therefore less subject to intra and interspecies variation in copy number. Further, the proportional assignment of reads to tree species showed good agreement between the DArTag and DArTseq platforms, providing confidence in the ability of these approaches to provide semi-quantitative information, despite the amplification step and scaling of the raw reads required with DArTag.

We have established that DArTag is a method of choice for identification of species-specific SNPs in koala faeces. The extremely low efficiency of species-specific SNPs detection by DArTseq (average of 20 reads in average 2.5 million sequencing depth, 0.0008%) combined with costly sequencing at depth makes this method unsuitable for determination of koala diet. With the DArTag assay we enriched the species-specific SNPs detection by 1,000 times on average (average of 432 reads in average 50,000 sequencing depth, 0.8%) at a fraction of the sequencing cost. Even with the pre-investment in oligo synthesis required for DArTag this method delivers much higher, albeit still low, efficiency at much lower cost. The work to increase the DArTag’s efficiency is already in progress. Better performance of DArTag is likely because this platform includes a hybridization and amplification step that enrich target DNA in the sample. This is of particular benefit in this system where dietary DNA comprises a very small fraction of the total DNA in the sample (unpublished data; Schultz et al., 2018).

The example applications of our method provided here have already shown that translocated koalas are able to readily accommodate previously unfamiliar eucalypt species into their diet, a finding with significant implications for koala management and conservation. Further, we have also revealed unexpected tree species to potentially be important components of the diets of some koalas. For instance, we found that some koalas in the Blue Mountains feed on *A. costata* and *C. gummifera,* with the former making up a significant proportion of some samples. While both tree species are known to be used by koalas for resting (OEH, 2018), both were strongly associated with sites not used by koalas in a previous study by Phillips and Callaghan (2000), and neither is generally viewed as a food tree (Cristescu, Banks, Carrick, & Frère, 2013). The ability of this method to identify previously overlooked food tree species is of course dependent upon these species being identified *a priori* as candidate food species and included in reference collections from the commencement of the project. In this respect, the method is at a potential disadvantage compared to DNA barcoding, which can in principle, draw upon extensive databases such as NCBI to identify food species. Nonetheless, where a suitable reference collection of potential food species is assembled this method is less costly, more time efficient and has high discrimination then other approaches used to date.

We anticipate that this method will be applied more intensively to koala individuals and populations to gain greater insight into diet composition in different habitats and across seasons. This information in turn will inform our understanding of koala habitat requirements and thereby direct management and conservation policy. We are also currently using this method to gain insight into the association between diet composition and the gastrointestinal microbiome. The development of custom SNP marker libraries also holds the distant promise of markers that can be associated with phenotypes of consumed species. As an example, koalas make feeding decisions not only among tree species, but also among individual trees, and these choices are strongly influenced by concentrations of plant secondary metabolites (Moore & Foley, 2005), which are under strong genetic control (Andrew, Wallis, Harwood, Henson, & Foley, 2007). At present, the genes responsible for these chemical defences in eucalypts are unknown, but the ability to non-invasively identify not only what species are consumed by a cryptic animal, but also the traits of the individuals consumed, is an exciting prospect.

## Acknowledgements

We thank Scott Bevins (since deceased) for his assistance in collecting koala faecal samples from the Blue Mountains and Aireys Inlet koalas. The following people assisted in collecting faecal samples from the Aireys Inlet koalas: Emily Scicluna, Kiarah Smith, Louise Falls, Linda Brown and Rachel Loneragan from the conservation Ecology Centre, Cape Otway; Emily Hynes and Martin Taube from Ecoplan, Australia. We thank Peter Menkhorst from the Arthur Rylah Institute for Environmental Research and the Department of Environment, Land, Water and Planning, Victoria for allowing us to undertake our research at Aireys Inlet in association with their koala translocation program. We acknowledge the traditional owners and their custodianship of the lands on which this study was conducted; the Gadabanud, Wathaurong and Darug people.

Fieldwork in the Blue Mountains was carried out under Western Sydney University Animal Ethics approval (A10670) and permits from the New South Wales Office of Environment and Heritage (SL101364 and SL100822). Fieldwork at Aireys Inlet was carried out under Western Sydney University Animal Ethics approval (A11253) and with an appropriate permit from the Victorian government (10007714).

The development of the genetic method for dietary determination was funded by grants from the Conservation Ecology Trust, as well as donations collected from the general public by WILD LIFE Sydney Zoo at Darling Harbour. Fieldwork at Aireys Inlet was funded by the Australian Research Council (LP140100751) in partnership with Evolva Biotech A/S, the NSW Office of Environment and Heritage and The Conservation Ecology Trust.

## Data Accessibility

The DArTseq raw sequencing reads for the eucalypt samples are available in NCBI under BioProject PRJNA791151.

## Author Contributions

MB contributed to the design of the project, collection of eucalypt and scat samples, DNA extraction, data analysis and writing the manuscript. KB collected the Mountain Lagoon scat samples, contributed to the DNA extractions and edited the manuscript. KHU conducted the bioinformatic analysis of the DArTseq and DArTag data, developed the laboratory protocols for the DArTag platform and contributed to writing the manuscript. JP collected scat samples from Airey’s Inlet and edited the manuscript. DJ designed the oligos for the DArTag platform, performed the bioinformatic analysis of the DArTag data and contributed to editing the manuscript. KL fitted koalas at Mountain Lagoon with VHF collars that allowed them to be located for sampling and edited the manuscript. BM contributed to the design of the project, collection of the eucalypt samples and writing the manuscript.

## Supplementary Tables

Table S1: Sampling locations of Blue Mountains Eucalyptus leaves

Table S2: Fix private alleles identified for the 22 potential dietary tree species from the Dartseq data

## Supplementary Figures

Fig S1: Frequency distribution for the proportion of reads that were the alternative allele. Only instances where both alleles where detected are shown.

Fig S2: Proportion of matches and mismatches between individual and pooled tree samples

Fig S3: The first two axes of the principal components analyse based on the Hamming distance matrix generated from DArTseq SNPs data for all the candidate koala food tree species at Mountain Lagoon and Aireys Inlet.

## Notes

### Competing Interest Statement

The authors have declared no competing interest.

