## Supplementary Figures for "A new genetic method for diet determination from faeces that provides species level resolution in the koala"

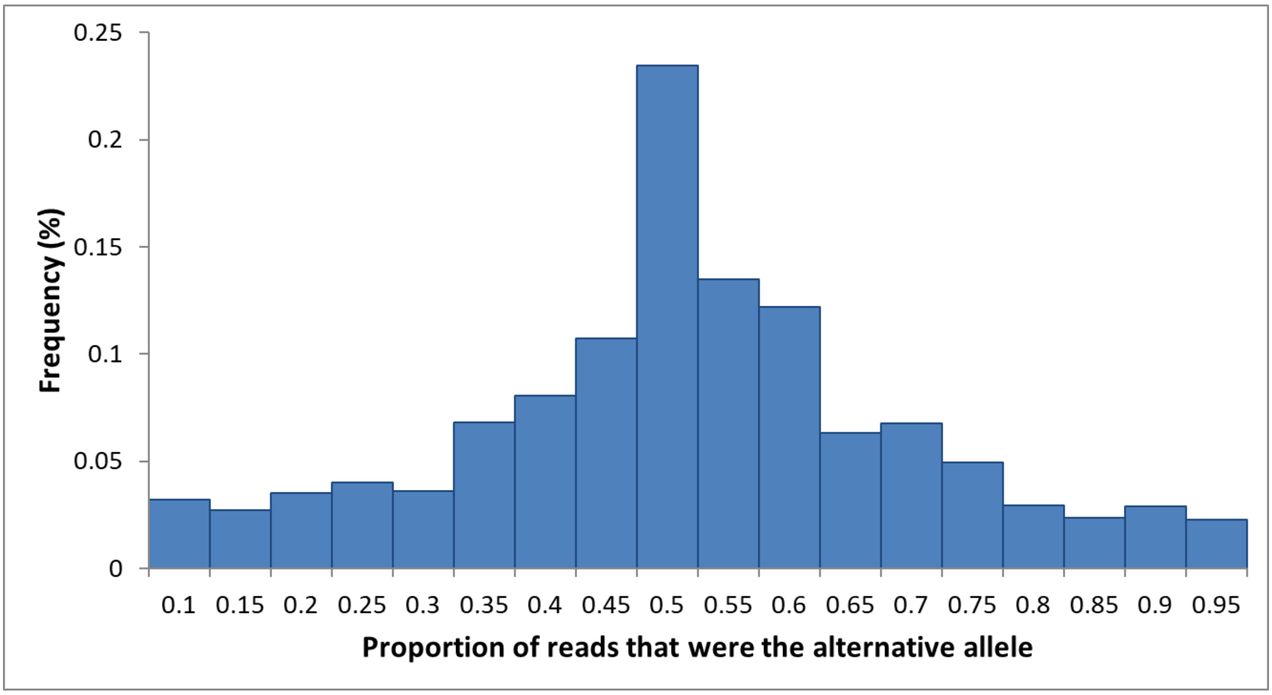


Fig S1: Frequency distribution for the proportion of reads that were the alternative allele. Only instances where both alleles where detected are shown.

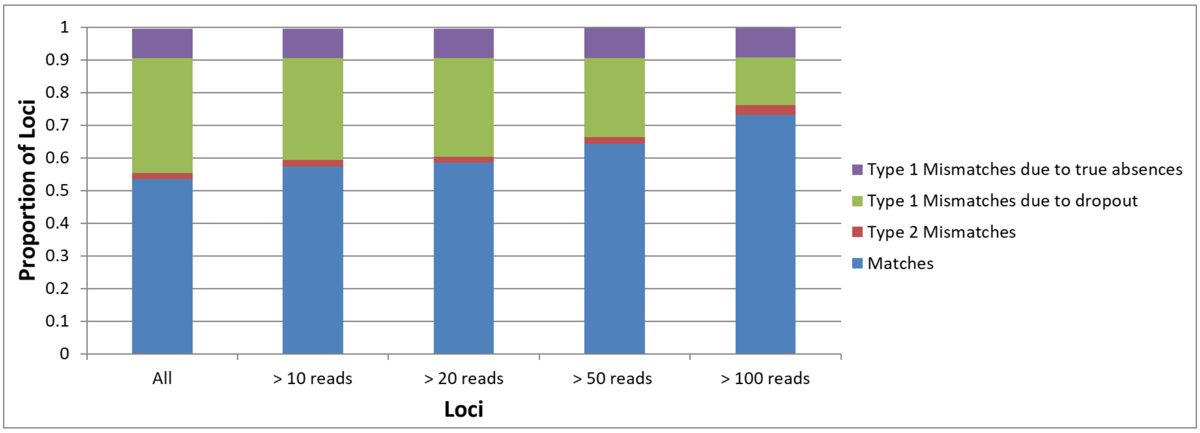


Fig S2: Proportion of matches and mismatches between individual and pooled tree samples

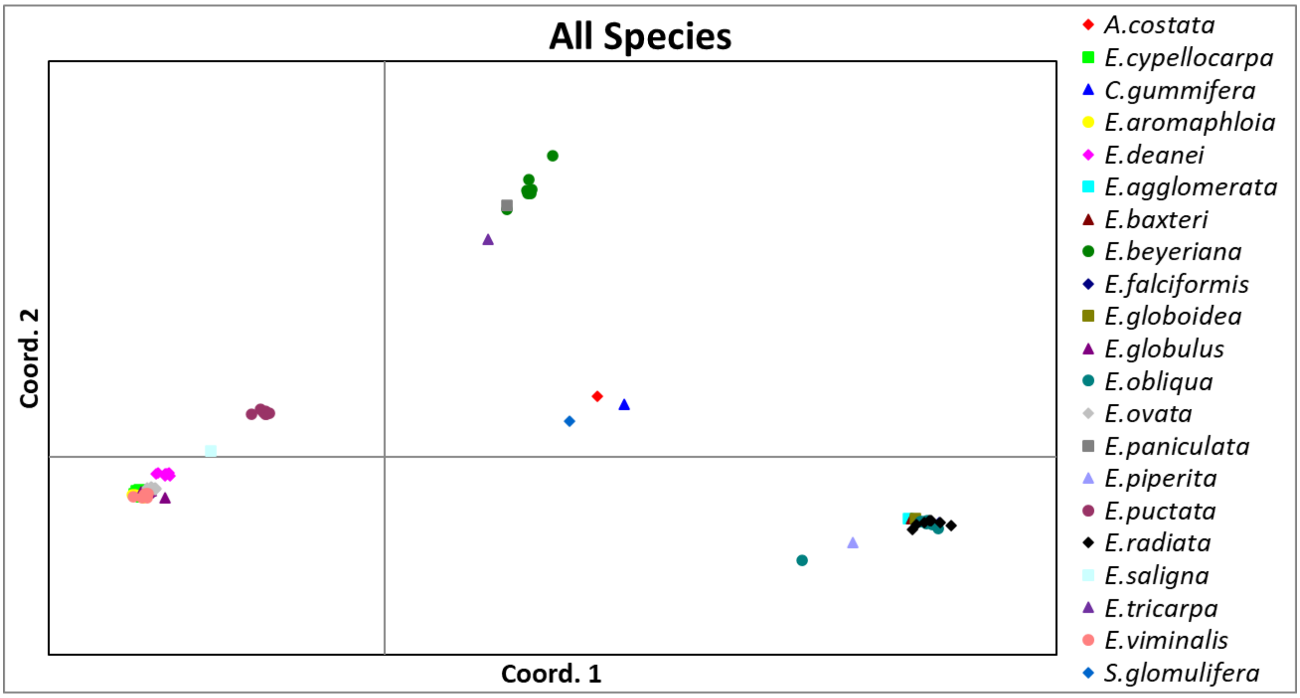


Fig S3: The first two axes of the principal components analyse based on the Hamming distance matrix generated from DArTseq SNPs data for all the candidate koala food tree species at Mountain Lagoon and Aireys Inlet.
